# The RNA polymerase gene specificity factor σ^54^ is required for homogeneous non-planktonic growth of uropathogenic *Escherichia coli*

**DOI:** 10.1101/2022.01.13.476216

**Authors:** Amy Switzer, Lynn Burchell, Panagiotis Mitsidis, Sivaramesh Wigneshweraraj

## Abstract

The canonical function of a bacterial sigma (σ) factor is to determine the gene specificity of the RNA polymerase (RNAP). In several diverse bacterial species, the σ^54^ factor uniquely confers distinct functional and regulatory properties on the RNAP. A hallmark feature of the σ^54^-RNAP is the obligatory requirement for an activator ATPase to allow transcription initiation. The genes that rely upon σ^54^ for their transcription have a wide range of different functions suggesting that the repertoire of functions performed by genes, directly or indirectly affected by σ^54^, is not yet exhaustive. By comparing the non-planktonic growth properties of prototypical enteropathogenic, uropathogenic and non-pathogenic *Escherichia coli* strains devoid of σ^54^, we uncovered σ^54^ as a determinant of homogenous non-planktonic growth specifically in the uropathogenic strain. Notably, bacteria devoid of individual activator ATPases of the σ^54^-RNAP do not phenocopy the σ^54^ mutant strain. It seems that σ^54^’s role as a determinant of homogenous non-planktonic growth represents a putative non-canonical function of σ^54^ in regulating genetic information flow.

## Introduction

Central to bacterial gene expression is transcription; the step where RNA synthesis occurs. Transcription in bacteria is catalysed by the multisubunit enzyme, RNA polymerase (RNAP). For promoter-specific initiation of transcription, the catalytic complex of the RNAP (α_2_ββ’ω) has to associate with one of the many different sigma (σ) factor subunits. The primary function of σ factors is to direct the RNAP to the promoters of specific sets of genes. Many bacterial genomes contain multiple σ factors, which reversibly and competitively bind to the RNAP to execute and coordinate specific transcription programmes in order to regulate cellular processes to a particular growth condition. As such, σ factors can be considered as master regulatory factors governing bacterial gene expression. Based on functional and structural criteria, bacterial σ factors are grouped into two distinct classes (1). In *Escherichia coli*, six of the seven σ factors belong to the σ^70^ class, named after the prototypical housekeeping σ factor, σ^70^. The σ^54^ factor, which is historically associated with executing transcriptional programmes that allow bacteria to cope with the conditions of nitrogen adversity (2) and found in genomes of many phylogenetically diverse bacteria, exists in a class of its own (3,4).

The σ^70^ and σ^54^ classes of σ factors confer distinct regulatory properties upon the RNAP. The σ^70^ class of σ factors (hereafter referred to as σ^70^ for simplicity) direct the RNAP to promoters with conserved sequences centered at -35 and -10 base pairs (bp) upstream from the transcription start site (+1 site). In contrast, σ^54^ directs its RNAP to promoters characterized by conserved sequences located at -24 and -12 bp upstream of the +1 site. The initial binding of the RNAP to the promoter results in a closed promoter complex (RPc). Regulation of transcription at most σ^70^-dependent promoters often occurs to either stimulate or antagonize RPc formation. The RPc at σ^70^-dependent promoters is usually short-lived and either dissociates or spontaneously isomerizes to the transcriptionally proficient open promoter complex (RPo). In the RPo, the promoter DNA strands are locally melted, such that the single-stranded +1 site is positioned in the catalytic cleft of the RNAP to allow initiation of RNA synthesis. The RPc at σ^54^-dependent promoters rests in a ‘ready-to-respond’ state for RPo formation. Conversion of the RPc to RPo at σ^54^-dependent promoters requires a specialized activator ATPase. Different activator ATPases sense and couple different environmental and intracellular signals to activate specific sets of σ^54^-dependent promoters. The activator ATPase binds to cognate DNA sites, called enhancers, located ∼100–150 bp frequently upstream but sometimes downstream of σ^54^-dependent promoters. The interaction between the enhancer-bound activator ATPase and the RPc occurs via a DNA-looping event and results in ATP hydrolysis-dependent conformational rearrangements in the σ^54^, the catalytic subunits of the RNAP, and the promoter DNA to allow RPo formation.

The reliance on different specialised activator ATPases allows the σ^54^-containing RNAP (σ^54^-RNAP) to rapidly activate specific transcription programmes to respond to a particular growth condition. As such, the products of genes that depend on σ^54^ have a wide range of different functions in *E. coli* and several related bacteria but no obvious single theme appears in the repertoire of functions performed by their products (5). Many animal and plant pathogenic bacteria rely on the σ^54^ for transcription of genes linked to virulence or virulence-associated processes (reviewed in (6)). *E. coli* is a commensal inhabitant of the mammalian gastrointestinal tract, but *E. coli* lineages have acquired specific virulence characteristics, which confer them the capacity to adapt and thrive in specific host niches, causing significant morbidity and mortality as human pathogens. Although pathogenic *E. coli* can cause intestinal/enteric and extra-intestinal infections – both with the potential to lead to serious systemic disease – most of our knowledge on the involvement of σ^54^ in *E. coli* pathogenesis is limited to studies on the intestinal pathogen, enterohemorrhagic *E. coli* (EHEC). To establish itself in the host, EHEC bacteria must overcome the acidic gastrointestinal environment and the σ^54^ is involved in the cascade of processes associated with conferring acid resistance and attachment to intestinal cells for competitive host colonization (6-9).

Uropathogenic *E. coli* (UPEC) are members of the extra-intestinal pathogenic *E. coli* and are the major causative agent of urinary tract infections (UTIs) worldwide. UTIs are among the most common bacterial infections in humans in community and nosocomial settings. UPEC infections range in severity and are a healthcare problem compounded by the emergence of antibiotic resistant UPEC isolates (reviewed in (10,11)). UPEC bacteria normally reside in the gastrointestinal tract but cause disease when they gain entry to the urinary tract leading to cystitis; the most common form of UTI. UPEC bacteria also invade the cytoplasm of uroepithelial cells, replicate and form intracellular bacterial communities (IBCs). Although the host immune system may remove some of the IBCs and excrete them with the urine, the remaining bacteria can exist as a biofilm resistant to host immune responses and antibiotic treatment. Some UPEC bacteria can escape from the biofilm and disseminate into the bladder lumen causing recurrent episodes of cystitis; some can ascend into the kidneys causing pyelonephritis; and some may also spread from the urinary tract to the bloodstream causing bacteraemia. Therefore, UPEC bacteria need to efficiently adjust their transcriptional programmes to adapt and survive in diverse host environments, resist assaults from the host’s immune system, and tolerate antibiotic therapy. Although the transcriptional programmes underpinning UPEC pathogenesis have been widely studied (e.g., (12-14)), little, if any, information exists on the role of σ^54^ in UPEC physiology and pathogenesis. We report a hitherto unknown role for σ^54^ in the UPEC strain CFT073, which was originally isolated from the blood and urine of a woman suffering from pyelonephritis (15).

## Results

### The planktonic and non-planktonic growth properties of different E. coli strains lacking σ^54^

We generated a Δ*rpoN* CFT073 strain and, intriguingly, when plated onto LB agar plates, we consistently observed two differently sized colonies following 48 h of incubation at 37 °C (Fig. 1*A*). In contrast, the wild-type CFT073 colonies were more homogeneous in size. The difference in colony size observed with the Δ*rpoN* CFT073 strain was reversible when *rpoN* was exogenously supplied via a low copy number plasmid from its native promoter (pACYC-*rpoN*). We did not detect any obvious difference in colony size when the Δ*rpoN* NCM3722 (a prototypic wild-type K-12 *E. coli* strain related to strain MG1655; (16)) or the Δ*rpoN* EDL933 (an EHEC strain) strains were plated onto LB agar plates, and the mutant colonies resembled those formed by the respective wild-type strains (Fig. 1*B* and 1*C*, respectively).

**Figure 1.**
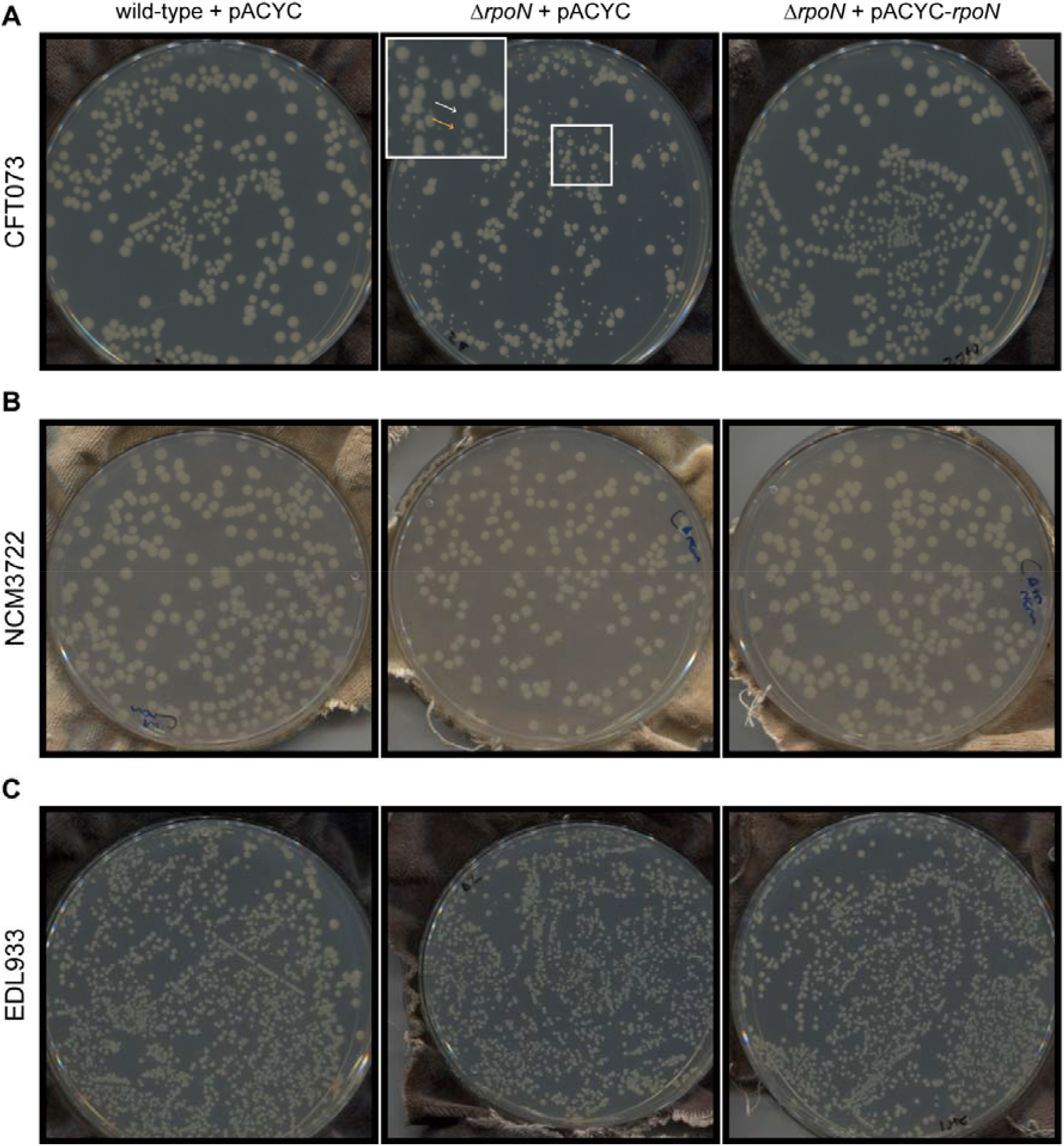
The non-planktonic growth properties of strains of *E. coli* lacking σ^**54**^. Images of colonies on LB agar plates incubated at 33 °C for 48 h containing wild-type (*left*), Δ*rpoN* (*middle*) or Δ*rpoN* + *rpoN* (*right*) of *E. coli* strains (*A*) CFT073, (*B*) NCM3722, and (*C*) EDL933. The two differently sized colonies seen on plates containing the Δ*rpoN* CFT073 strain are indicated with white (big colony) and orange (small colony) arrows in the inset in (*A*).

To better understand how σ^54^ contributes to the growth dynamics of the three different *E. coli* strains on LB agar plates, particularly the CFT073 strain, we used a method called ScanLag (17). This involves putting the inoculated LB agar plates onto a conventional office scanner, which itself is placed in a 33°C incubator, and periodically (in this case every 20 min) activating the scanner to measure the appearance time (in hours; h) of individual colonies and their apparent growth rate (in pixels^2^/h; px^2^/h; calculated during the first six hours after initial appearance). To inoculate the bacteria onto LB agar plates, we sub-cultured overnight LB liquid cultures of the wild-type, Δ*rpoN* and Δ*rpoN* + pACYC-*rpoN* bacteria into fresh LB liquid media and grew the bacteria for 5 h at 37°C. At this stage, the bacteria were in the early stationary phase of growth (see below), and we plated 100μl of each culture diluted by approximately 10^−5^-10^−6^ for the ScanLag experiment. For the Δ*rpoN* CFT073 strain, we initially used a ‘big’ colony (see Fig. 1*A*) for the ScanLag experiments. The wild-type CFT037 strain had a doubling time of ∼0.66 h in LB liquid media (Fig. 2*A*) and its colonies appeared ∼10 h after incubation and had an apparent growth rate of ∼33 px^2^/h (Fig. 2*B*). The ‘big’ colony of the Δ*rpoN* CFT073 strain had a slightly longer doubling time in LB liquid media compared to the wild-type strain (Fig. 2*A*). Notably, the appearance time of about half the proportion of the Δ*rpoN* CFT073 colonies (hereafter referred to as M1) was delayed by ∼5 h compared to the wild-type strain and had an apparent growth rate of ∼22 px^2^/h, while the appearance time of the other proportion of the Δ*rpoN* CFT073 colonies (hereafter referred to as M2) was delayed by ∼20 h compared to the wild-type strain and had an apparent growth rate of ∼6 px^2^/h (Fig. 2*B*). The colonies of the Δ*rpoN* CFT073 bacteria containing plasmid pACYC-*rpoN*, which exogenously produces σ^54^ from its native promoter, as expected, neither displayed a growth defect in LB liquid media (Fig. 2*A*) nor the biphasic colony appearance pattern (Fig. 2*B*). We then picked the M1 and M2 colonies from the LB agar plate, created an overnight inoculum in LB liquid media, and compared their growth properties in fresh LB liquid media and in ScanLag experiments. As shown in Fig. 2*C*, the doubling time of the bacteria from the M1 and M2 colonies did not differ, suggesting that the slow apparent growth rate of the M2 colony on LB solid media is reversible in LB liquid media. Consistent with this view, in ScanLag experiments, bacteria grown from M1 and M2 colonies displayed an indistinguishable biphasic appearance pattern on LB agar plates (Fig. 2*D*). Although the doubling time of the Δ*rpoN* NCM3722 was also compromised in LB liquid media (like the Δ*rpoN* CFT073 strain) (Fig. 2*E*), the Δ*rpoN* NCM3722 strain, consistent with results in Fig. 1*B*, did not display a biphasic appearance pattern on LB agar plates (Fig. 2*F*). However, the appearance time of Δ*rpoN* NCM3722 colonies on LB agar plates was delayed by ∼7 h (apparent growth rate ∼20 px^2^/h) compared to the wild-type NCM3722 colonies (apparent growth rate ∼63 px^2^/h) (Fig. 2*F*). The Δ*rpoN* EDL933 strain neither displayed any growth defect in LB liquid media (Fig. 2*G*) nor the biphasic appearance pattern on LB agar plates (Fig. 2*H*). However, the appearance time of Δ*rpoN* EDL933 colonies on LB agar plates was delayed by ∼5 h (apparent growth rate ∼18 px^2^/h) compared to the wild-type EDL933 colonies (apparent growth rate ∼19 px^2^/h) (Fig. 2*G*). As with the Δ*rpoN* CFT073 strain, the presence of plasmid pACYC-*rpoN* reverted the LB liquid growth defect (where applicable) and returned colony appearance time of the Δ*rpoN* NCM3722 and Δ*rpoN* EDL933 strains to their respective wild-type levels. Overall, (1) the absence of σ^54^ in *E. coli* strains NCM3722, EDL933 and CFT073 results in a notably delayed appearance of colonies on LB agar plates, despite only having a modest adverse effect on growth in LB liquid media; (2) in the case of the *E. coli* strain CFT073, the absence of σ^54^ significantly delays the appearance time of a subpopulation of bacteria on LB agar plates (i.e., the M2 colonies); and (3) the delayed appearance of this subpopulation of Δ*rpoN* CFT073 colonies is a property specific to the CFT073 strain as the biphasic colony appearance pattern is not obviously apparent with the NCM3722 and EDL933 strains. In conclusion, it seems that σ^54^ is required for homogeneous non-planktonic growth of *E. coli* strain CFT073.

**Figure 2.**
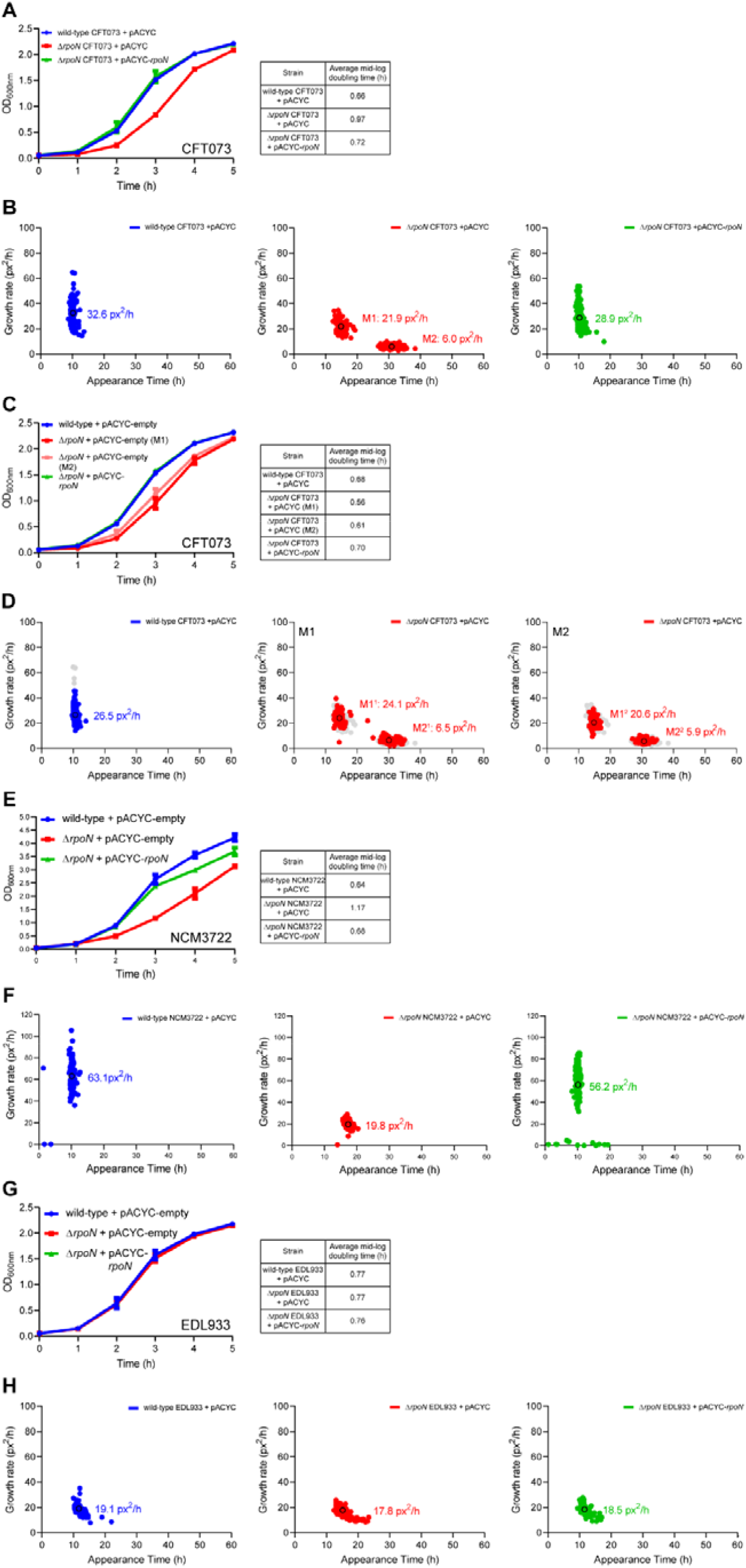
The planktonic and non-planktonic growth properties of different strains of *E. coli* lacking σ^54^. *A*, Graph showing the growth curves of wild-type (blue), Δ*rpoN* (red), and Δ*rpoN* + *rpoN* (green) CFT073 strains grown in LB liquid media for 5 h. *B*, Scanlag analysis of colony appearance time for CFT073 strains grown in LB liquid media for 5 h of wild-type (*left*), Δ*rpoN* (*red*), and Δ*rpoN* + *rpoN* (*green*). Black circles represent average population growth rate and appearance time with mean growth rate indicated as a value. *C*, Graph showing the growth curves in LB liquid media from colonies picked from (*B*) and colour coded as indicated. The average doubling time for each colony type is indicated in the table. *D*, Scanlag analysis as in (*B*) for bacteria plated from (*C*). Shown in grey is data from (*B*) for reference. *E*, Graph showing the growth curves of wild-type (*blue*), Δ*rpoN* (*red*), and Δ*rpoN* + *rpoN* (*green*) NCM3722 strains grown in LB liquid media for 5 h. *F*, Scanlag analysis of colony appearance time as in (*B*) for NCM3722 strains. *G*, Graph showing the growth curves of wild-type (*blue*), Δ*rpoN* (*red*), and Δ*rpoN* + *rpoN* (*green*) EDL933 strains grown in LB liquid media for 5 h. *H*, Scanlag analysis of colony appearance time as in (*B*) for EDL933 strains grown in LB liquid media for 5 h. Where indicated, the error bars represent standard deviation (n□= □3).

### Heterogeneous non-planktonic growth is an inherent property of ΔrpoN CFT073 bacteria

We wanted to determine how the appearance time of colonies M1 and M2 of the Δ*rpoN* CFT073 strain are affected by the growth stage and culture conditions of the bacteria used for the ScanLag experiments. The colony appearance time and pattern of stationary phase bacteria on LB agar plates (Fig. 2*B*) served as a reference condition (hereon shown in grey in all applicable figures). As shown in Fig. 3*A*, when exponentially growing bacteria were used for the ScanLag experiment, both M1 and M2 colonies were present. However, we note that the M2 colonies of exponentially growing bacteria appeared sooner (by ∼5 h) than the M2 colonies of early stationary phase bacteria. Similarly, when bacteria from a 5-day old culture were used, the M2 colonies were still present but they appeared sooner (by ∼8 h) than the M2 colonies of early stationary phase bacteria (Fig. 3*B*). We next wanted to determine how bacteria grown in healthy female urine (HFU) affected the appearance time of colonies M1 and M2 on LB agar plates. As shown in Fig. 3*C*, the doubling time of the Δ*rpoN* CFT073 strain in HFU did not substantially differ from that of the wild-type CFT073 strain. Although, unsurprisingly, the doubling time of all strains was markedly slower in HFU than in LB liquid media (Fig. 3*C* and Fig. 2*A*, respectively). In the ScanLag experiments, the biphasic colony appearance pattern of Δ*rpoN* CFT073 colonies was evident, but the apparent growth rates of both M1 and M2 colonies were reduced compared to those of Δ*rpoN* CFT073 grown in LB (Fig. 3*D* and Fig. 2*B*, respectively). We also conducted ScanLag experiments with HFU-grown bacteria plated onto agar plates containing 50% (v/v) of HFU. In this case, the apparent growth rates of colonies of wild-type, Δ*rpoN* and Δ*rpoN* containing plasmid pACYC-*rpoN* CFT073 strains were reduced (Figure 3*E*). Although the colonies of the Δ*rpoN* strain appeared in a biphasic fashion, we note a marked change in the timing of appearance of the M2 colonies, which appeared sooner (by ∼5 h) than the M2 colonies of the HFU-grown Δ*rpoN* bacteria plated onto normal LB agar plates (Fig. 3*D* and 3*E*). We also note the presence of a small population of wild-type colonies that appear ∼37-40 h after incubation, but like in other conditions, most of the wild-type colonies appeared ∼10 h after incubation. In all conditions tested, as expected, the appearance time of colonies of the Δ*rpoN* CFT073 bacteria containing plasmid pACYC-*rpoN* resembled that of the wild-type strain. Overall, it seems that heterogeneous non-planktonic growth is an inherent property of the CFT073 strain when *rpoN* is absent and that the slower growing subpopulation of colonies (i.e., the M2 colonies) appear to be more sensitive to the growth environment and conditions than the faster growing colonies (i.e., the M1 colonies). In conclusion, the results underscore that σ^54^ is a determinant for the homogeneous non-planktonic growth of *E. coli* CFT073.

**Figure 3.**
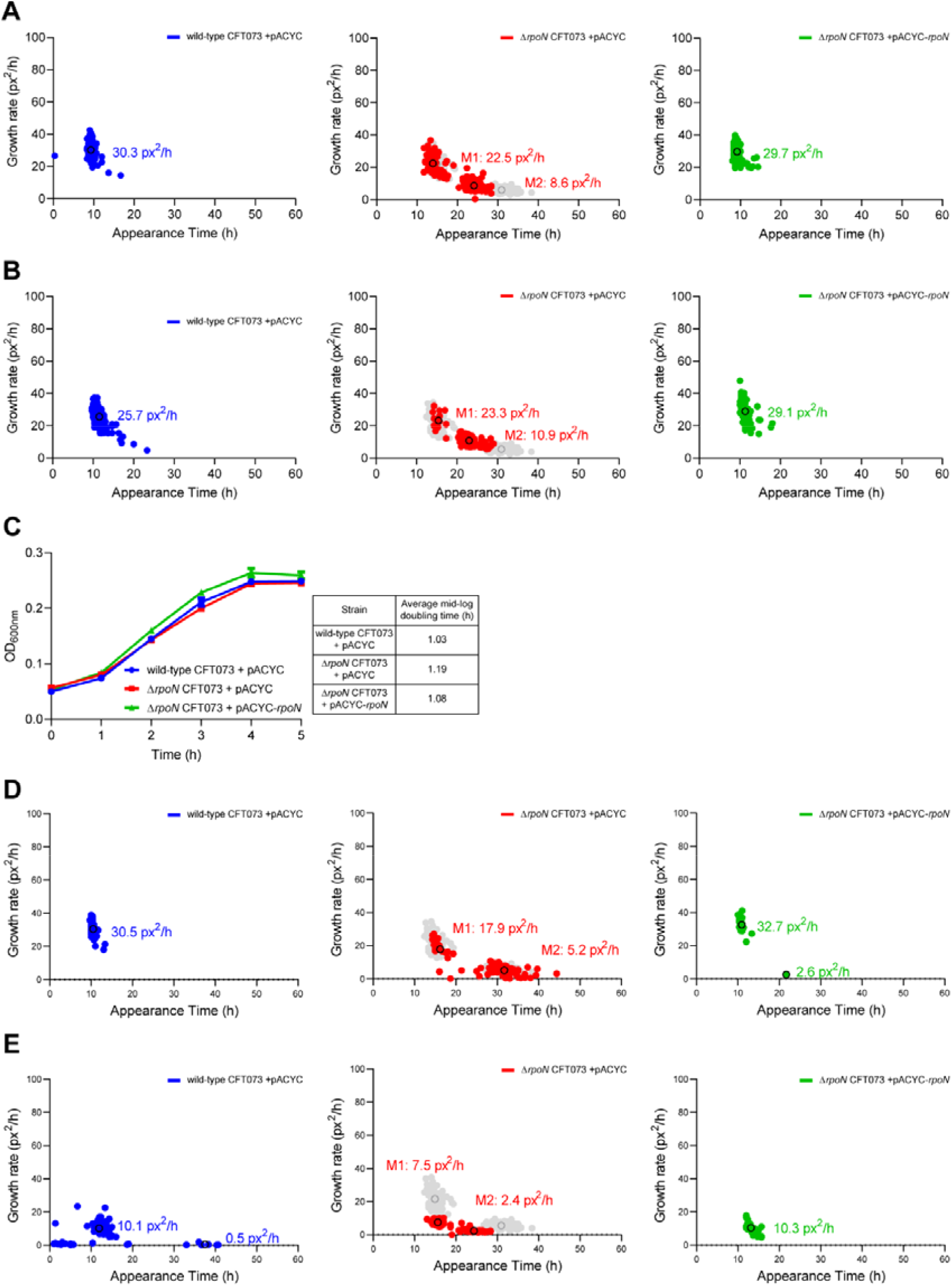
Heterogeneous non-planktonic growth is an inherent property of Δ*rpoN* CFT073 bacteria. *A*, Scanlag analysis of colony appearance time for wild-type (left, blue), Δ*rpoN* (middle, red), and Δ*rpoN* + *rpoN* (right, green) CFT073 bacteria grown in LB liquid media to mid-exponential phase before plating on LB agar plates. Black circles represent average population growth rate and appearance time with mean growth rate indicated as a value. Shown in grey for comparison are the M1 and M2 colonies from bacteria grown in LB liquid media for 5 h before plating. *B*, As in (*A*) but with bacteria grown for 5 days in LB liquid media. *C*, Graphs showing growth curves of wild-type (*blue*), Δ*rpoN* (*red*), and Δ*rpoN* + *rpoN* (*green*) CFT073 strains grown in human female urine for 5 h. *D*, As in (*A*) but with bacteria from (C). *E*, As in (*D*) but bacteria were plated onto agar plates containing 50% (v/v) human female urine. Where indicated, the error bars represent standard deviation (n□= □3).

### Comparative analysis of the transcriptomes of wild-type and ΔrpoN CFT073 strains

To better understand the requirement for σ^54^ for homogeneous non-planktonic growth of *E. coli* CFT073, we compared the transcriptomes of exponentially growing wild-type, Δ*rpoN* and Δ*rpoN* bacterial containing plasmid pACYC-*rpoN*. We defined differentially expressed genes as those with expression levels changed ≥2-fold with a false discovery rate-adjusted *P*-value <0.05. A total of 159 genes were differentially expressed in the Δ*rpoN* CFT073 strain (Table S2 and Fig. 4*A*), of which 149 (2.7 % of total genes) were upregulated and 10 (0.2 % of total genes) were downregulated. Of the genes that were downregulated, *glnHPQ, ybeJ* and *pspA* are transcribed from known σ^54^-dependent promoters. The products of *glnHPQ, ybeJ*, and *pspA* are involved in nitrogen and membrane stress responses, respectively, and their transcription is dependent on the activator ATPases, NtrC and PspF, respectively. To the best of our knowledge, *E. coli* cells growing exponentially in LB liquid media do not experience any nitrogen limitation or membrane stress. Therefore, it is unlikely that the downregulation of *glnHPQ, ybeJ* and/or *pspA* underpins the biphasic appearance of Δ*rpoN* CFT073 colonies on LB agar plates (also see later). The unknown gene product, c0897, is categorized as significantly downregulated, but genomic visualization revealed that this was due to its overlap with the *glnHPQ* promoter, and the gene product itself is not differentially expressed, so has been omitted from downstream analysis. We next focused on the genes that are upregulated in the Δ*rpoN* CFT073 strain. As shown in Fig. 4*B*, we noted that the majority (76 out of 149) of the conserved upregulated genes in the Δ*rpoN* CFT073 strain either belonged to or are associated with the *rpoS* regulon (as defined in the non-pathogenic *E. coli* strain MG1655 (18,19)) and transcribed by the general stress response σ factor, σ^38^ (product of *rpoS*). The functional interconnection between the *rpoN* and *rpoS* regulons is undisputed as the absence of *rpoN* has been shown to lead to either increased σ^38^ levels or σ^38^ stability in both enterohemorrhagic and non-pathogenic *E. coli* strains (7,8,20). Therefore, we considered whether the biphasic non-planktonic growth property of the Δ*rpoN* CFT073 strain was linked to increased σ^38^ activity. To investigate this, we constructed Δ*rpoS* and Δ*rpoN*Δ*rpoS* CFT073 strains and compared their growth properties in LB liquid media and by Scanlag analysis. As shown in Fig. 4*C*, the doubling time of the Δ*rpoN*Δ*rpoS* CFT073 strain did not differ from that of the Δ*rpoN* CFT073 strain. Further, the doubling time of the Δ*rpoS* CFT073 strain did not differ from that of the wild-type CFT073 strain. Notably, in the Scanlag experiment, the colonies of the Δ*rpoN*Δ*rpoS* CFT073 strain appeared in a biphasic fashion and their apparent growth rates and appearance times resembled that of those seen with the Δ*rpoN* CFT073 strain (Fig. 4*D*). However, we note that the M2 colonies of the Δ*rpoN*Δ*rpoS* CFT073 strain appeared faster (by ∼ 4 h) than the M2 colonies of the Δ*rpoN* CFT073 strain. As expected, the colonies of Δ*rpoN*Δ*rpoS* CFT073 strain containing plasmid pACYC-*rpoN* resembled wild-type colonies (Fig. 4*E*). Further, the colonies of the Δ*rpoS* CFT073 strain did not grow in a biphasic fashion and resembled wild-type colonies (Fig. 4*F*). Overall, it seems that *rpoS* does not directly contribute to the biphasic non-planktonic growth property of the Δ*rpoN* CFT073 strain. As many of the derepressed genes have no assigned function (Fig. 4*G*), at this stage in our analysis, it is difficult to delineate the genetic pathways that contribute to the biphasic non-planktonic growth property of the Δ*rpoN* CFT073 strain in greater granularity. However, as shown in Fig. 2*H*, the Δ*rpoN* EDL933 strain does not display the biphasic non-planktonic growth property. Therefore, we compared the transcriptomes of exponentially growing wild-type, Δ*rpoN* and Δ*rpoN* + pACYC-*rpoN* EDL933 bacteria and observed that a total of 41 genes were differentially expressed in the Δ*rpoN* EDL933 strain (Table S3 and Fig. 4*H*), of which 15 (0.3% of total genes) were upregulated and 26 (0.5% of total genes) were downregulated. Most of the upregulated genes were associated with amino acid metabolism (histidine and threonine) and many of the genes downregulated in the Δ*rpoN* CFT073 and EDL933 strains were the same (i.e., *glnH, glnP, glnQ, ybeJ* and *pspA*). However, of the 15 differentially expressed genes in the Δ*rpoN* EDL933 strain only 1 was commonly upregulated between the two Δ*rpoN* mutant strains (Fig. 4*H -inset*) – *gadC*, which encodes an antiporter as part of the glutamate-dependent acid resistance system. Notably, while many of the σ^38^-dependent acid resistance system genes are upregulated in the Δ*rpoN* CFT073 strain, no other associated genes are similarly upregulated in the Δ*rpoN* EDL933 strain. Overall, it seems that, under our experimental conditions, where σ^54^-dependent transcription is not known to be essential, the absence of *rpoN* in CFT073 and EDL933 *E. coli* strains leads to de-repression of different sets of genes. Hence, it seems that, under conditions where σ^54^-dependent transcription is not essential, σ^54^ has a non-canonical regulatory function in transcriptionally repressing gene expression. In the case of the CFT073 *E. coli*, this potential non-canonical regulatory function is associated with promoting homogeneous non-planktonic growth of CFT073 bacteria.

**Figure 4.**
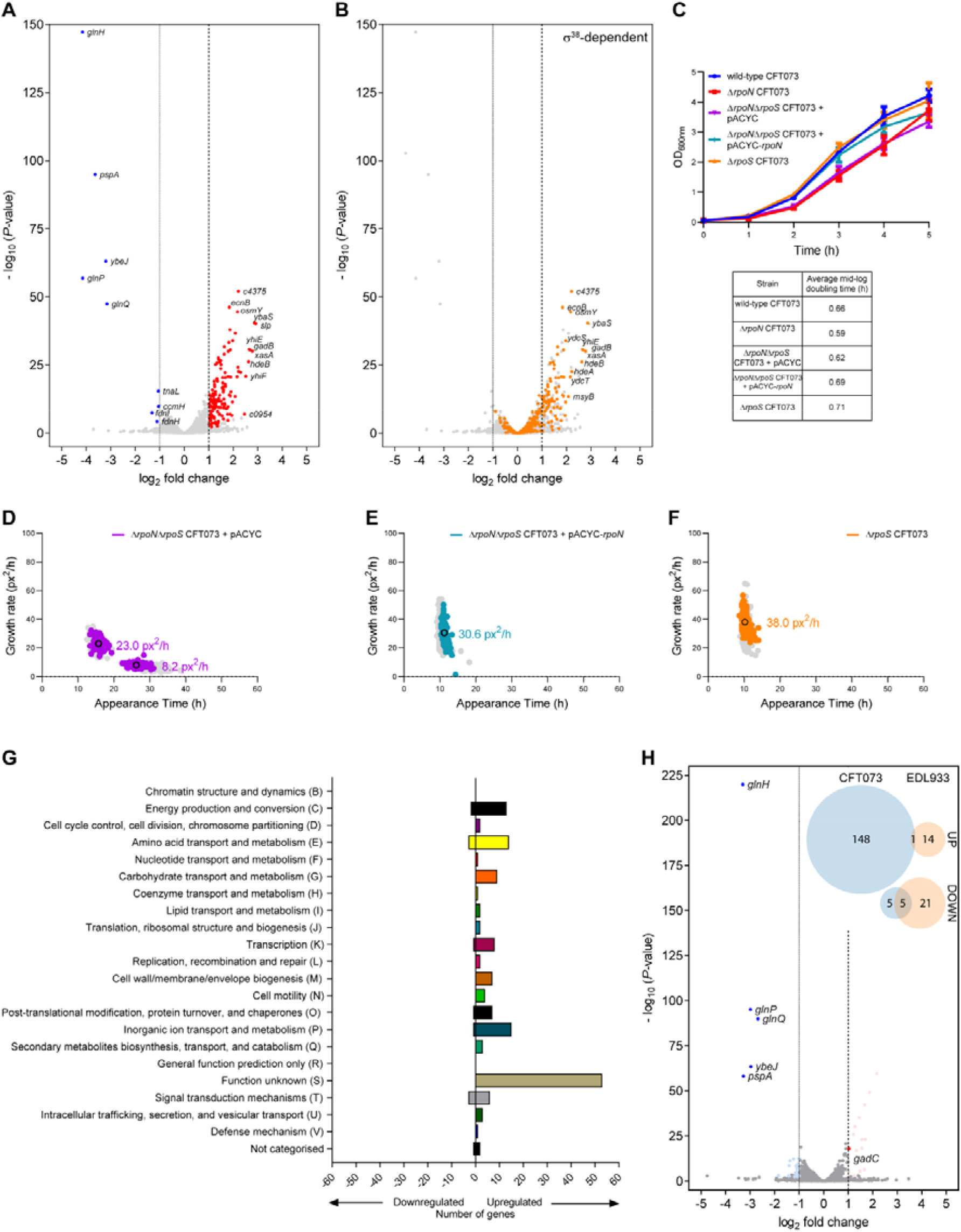
Comparative analysis of the transcriptomes of wild-type and Δ*rpoN* CFT073 strains. *A*, Volcano plot showing differentially expressed genes in Δ*rpoN* CFT073 strain as a log_2_ fold change from wild-type CFT073 strain extracted from LB liquid media during mid-exponential growth. Significantly differentially expressed genes (DEGs) were defined as having an absolute log_2_ fold change ≥ 1, and a false discovery rate-adjusted *P* value < 0.05. Upregulated DEGs shown in red, downregulated DEGs show in blue, and largest log_2_ fold changes labelled with gene names. *B*, As in (*A*) but known σ^38^-dependent genes highlighted in orange. Gene names of σ^38^-dependent genes with the highest log_2_ fold change are labelled. *C*, Graph showing growth curves of wild-type (*blue*), Δ*rpoN* (*red*), Δ*rpoN*Δ*rpoS* (*purple*), Δ*rpoN*Δ*rpoS* + *rpoN* (*turquoise*), and Δ*rpoS* (*orange*) CFT073 strains grown in LB liquid media for 5 h. Error bars represent standard deviation (n□= □3). *D*-*F*, Scanlag analysis of colony appearance time of (*D*) Δ*rpoN*Δ*rpoS* (*purple*), (*E*) Δ*rpoN*Δ*rpoS* + *rpoN* (*turquoise*), and (*F*) Δ*rpoS* (*orange*) CFT073 strains grown in LB liquid media for 5 h before plating. Black circles represent average population growth rate and appearance time with mean growth rate indicated as a value. Shown in grey for comparison are the M1 and M2 colonies from bacteria grown in LB liquid media for 5 h before plating. *G*, Graph showing DEGs in the Δ*rpoN* CFT073 bacteria from (*A*) categorised by COG annotation. *H*, As in (*A*) but for DEGs in Δ*rpoN* EDL933 bacteria as a log_2_ fold change from wild-type EDL933 bacteria extracted from LB liquid media during mid-exponential growth. Upregulated DEGs shown in pale red and downregulated DEGs shown in pale blue, with DEGs also present in (*A*) labelled and shown in bright red and bright blue, respectively. The Venn diagrams (*inset*) show upregulated and downregulated DEG numbers from Δ*rpoN* CFT073 bacteria (*blue*) compared to Δ*rpoN* EDL933 bacteria (*orange*) with numbers in overlapping circles representing identical genes in both strains.

### Activator ATPases do not appear to contribute to σ^54^’s role as a determinant for the homogeneous non-planktonic growth of E. coli CFT073

To further establish the potential non-canonical regulatory function of σ^54^, we focused on the activator ATPases. Recall that the σ^54^-RNAP requires specialized activator ATPases for its canonical function as a promoter-specificity factor. Our rational was that, if the biphasic non-canonical growth property of the Δ*rpoN* CFT073 strain was due to a canonical function of σ^54^ then a CFT073 strain lacking any one of the activator ATPases would also phenocopy the Δ*rpoN* CFT073 strain in the ScanLag assay. Conversely, this would be not the case if a non-canonical regulatory function of σ^54^ was responsible for the biphasic non-planktonic growth property of the Δ*rpoN* CFT073 strain. Hence, we searched the CFT073 genome for proteins that have the hallmark ‘GAFTGA’ sequences of activator ATPases that is required for interacting with the σ^54^-RNAP and identified 11 activator ATPases. As shown in Fig. 5 (inset graphs), the 11 activator ATPase mutant strains of CFT073 can be classified into two classes based on their growth properties in LB liquid media in a plate reader incubator at 37°C: The Δ*atoC*, Δ*yfhA*, Δ*prpR*, Δ*glnG* and Δ*c5040* displayed markedly compromised growth compared to the wild-type, Δ*rpoN* and other activator ATPase mutant strains. Thus, for the ScanLag experiment, we let each activator ATPase mutant strain reach early exponential phase of growth and based on optical density correction, plated approximately equal numbers of bacteria onto LB agar plates. The results showed that, for each activator ATPase mutant CFT073 strain, the differences in growth observed in LB liquid media were reflected in the apparent growth rate of the colonies on LB agar plates (Fig. 5). However, we did not detect the characteristic biphasic appearance of colonies, as seen with the Δ*rpoN* CFT073 strain, for any of the activator ATPase mutant CFT073 strains. We note that some Δ*yfhA* colonies resembled the M2 colonies of the Δ*rpoN* strain but, unlike the M2 colonies of the Δ*rpoN* strain, the frequency of the M2 colonies of the Δ*yfhA* strain represented <10% of the total number of colonies on the plate. As activator ATPases confer narrow regulon specificity upon the σ^54^-RNAP, it is unlikely that more than one activator ATPases are involved in σ^54^’s role as a determinant for the homogeneous non-planktonic growth of *E. coli* CFT073. Overall, our results indicate that none of the absence of any one of the activator ATPases encoded by the CFT073 strain phenocopy the Δ*rpoN* CFT073 strain, further supporting the view that the biphasic non-planktonic growth phenotype of the Δ*rpoN* CFT073 strain is due to a potential non-canonical regulatory function of σ^54^. in the CFT073 strain under the conditions used here.

**Figure 5.**
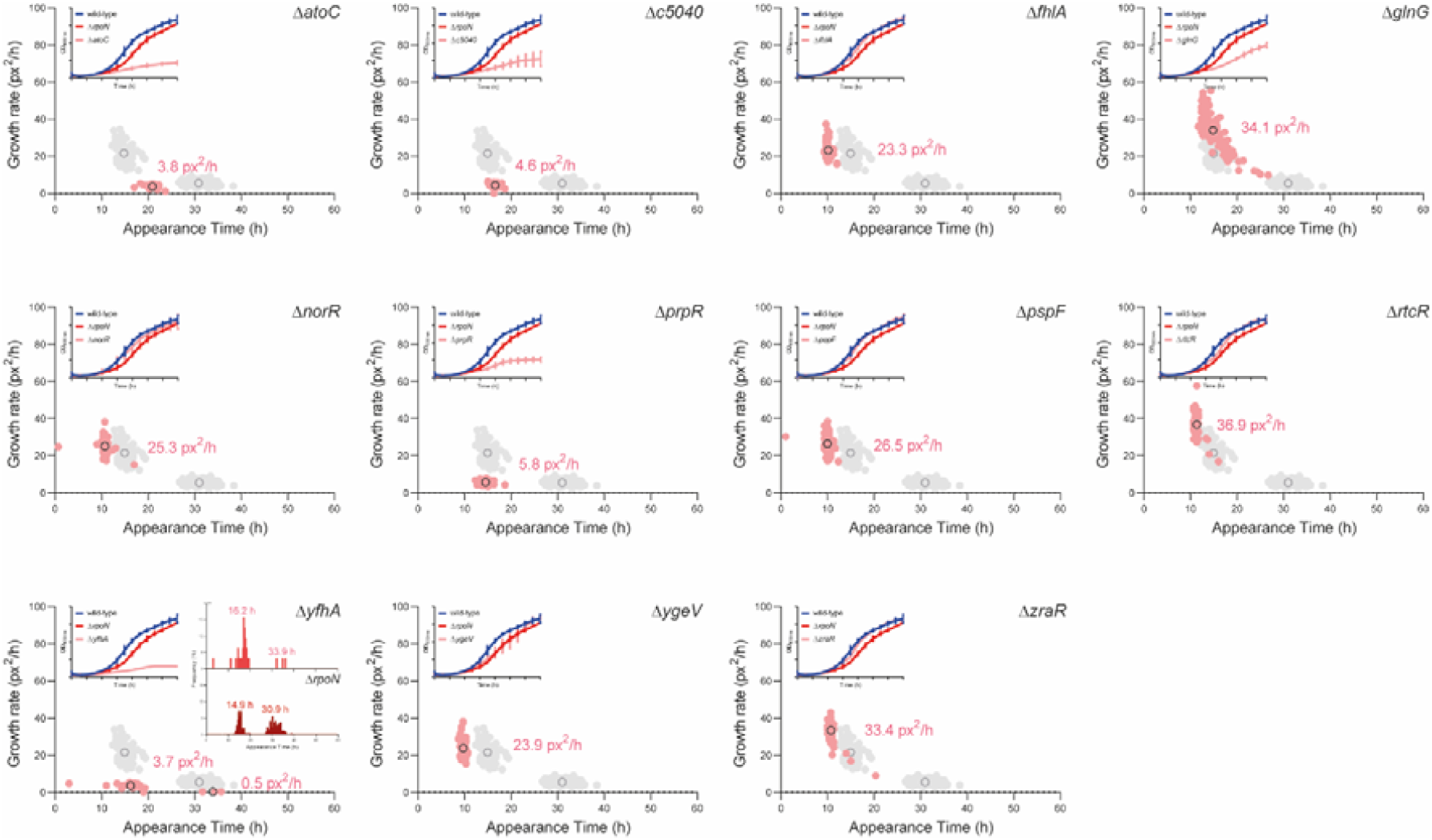
Activator ATPases do not appear to contribute to σ^54^’s role as a determinant for the homogeneous non-planktonic growth of *E. coli* CFT073. Scanlag analysis of colony appearance time following growth in LB liquid media for 5 h of known activator ATPase mutants of CFT073 bacteria in order from top left: Δ*atoC*, Δ*c5040*, Δ*fhlA*, Δ*glnG*, Δ*norR*, Δ*prpR*, Δ*pspF*, Δ*rtcR*, Δ*yfhA*, Δ*ygeV*, and Δ*zraR*. Black circles represent average population growth rate and appearance time with mean growth rate indicated as a value. Shown in grey for comparison are the M1 and M2 colonies from bacteria grown in LB liquid media for 5 h before plating. The inset graphs show growth curves of each activator ATPase mutant strain (*pink*) grown in LB for 5h with growth curves of the wild-type (*blue*) and Δ*rpoN* (*red*) CFT073 strains shown for comparison. The inset histograms for the Δ*yfhA* CFT073 strain show appearance times of Δ*yfhA* (*top, pink*) and Δ*rpoN* (*bottom, red*) for comparison (see text) with average population appearance time written above each peak. Where indicated, the error bars represent standard deviation (n□= □3).

### Fitness advantage conferred by homogeneous non-planktonic growth

The results thus far unambiguously indicate a potential non-canonical role for σ^54^ in ensuring homogeneous non-planktonic growth specifically of CFT073 bacteria. As it is widely accepted that growth heterogeneity can confer considerable fitness benefits to a bacterial population as a whole, we considered whether homogeneous non-planktonic growth has a fitness advantage under growth-restrictive conditions. As UPEC bacteria are frequently exposed to antibiotics, we investigated the sensitivity of M1 and M2 colonies of Δ*rpoN* CFT073 bacteria to minimum inhibitory concentration (MIC) of amikacin and nitrofurantoin – two frequently used antibiotics to treat UTIs caused by UPEC – against wild-type bacteria. We did this by conducting ScanLag analysis using LB agar plates containing MICs of amikacin or nitrofurantoin. As shown in Fig. 6*A*, we failed to detect any M2 colonies on LB agar plates containing MIC of amikacin; even though the appearance time of M1 colonies was like that of bacteria on LB agar plates without any antibiotics, they grew considerably slower on antibiotic-containing plates. Consistent with this observation, viability of Δ*rpoN* CFT073 bacteria was significantly reduced compared with wild-type CFT073 bacteria on LB agar plates with MIC of amikacin, suggesting that, rather than reverting to an M1 phenotype, the M2 population is killed in the presence of MIC levels of amikacin. In contrast, we detected the characteristic biphasic growth property of the Δ*rpoN* CFT073 bacteria on LB agar plates containing nitrofurantoin. However, the appearance time and apparent growth rate of both M1 and M2 colonies were markedly compromised compared to their growth on LB agar plates without any antibiotics, to such a degree that the M2 colonies were barely detectable (Fig. 6*B*). However, the overall viability of Δ*rpoN* CFT073 bacteria was comparable to wild-type CFT073 bacteria. We note that the appearance time and apparent growth rate of wild-type CFT073 bacteria on nitrofurantoin-containing LB agar plates are also compromised but not to the degree seen with the Δ*rpoN* CFT073 bacteria. Collectively, it seems that homogeneous non-planktonic growth, which requires *rpoN*, is likely to confer fitness advantages to the *E. coli* CFT073 strain under growth restrictive conditions.

**Figure 6.**
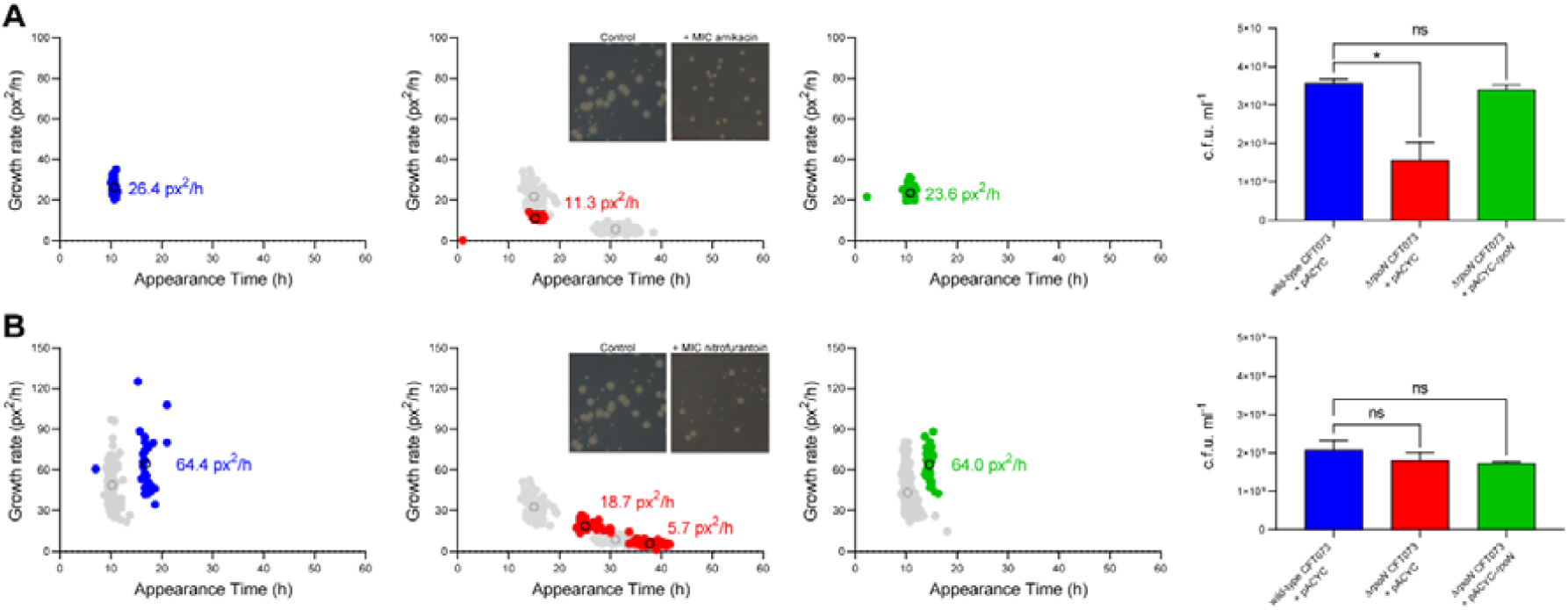
Fitness advantage conferred by homogeneous non-planktonic growth. *A*, Scanlag analysis of colony appearance time of wild-type (blue), Δ*rpoN* (red), and Δ*rpoN* + *rpoN* (green) CFT073 bacteria following growth in LB liquid media for 5 h followed by plating on LB agar plates with MIC of amikacin. Black circles represent average population growth rate and appearance time with mean growth rate indicated as a value. Shown in grey for comparison are the M1 and M2 colonies from bacteria grown in LB liquid media for 5 h before plating on LB agar containing no antibiotics. *B*, As in (*A*) but bacteria were plated on LB agar plates containing MIC of nitrofurantoin. The inset images show representative colonies for comparison (see text). In (*A*) and (*B*), the bar graphs indicate viability as measured by colony forming units (c.f.u.) of wild-type (*blue*), Δ*rpoN* (*red*), and Δ*rpoN* + *rpoN* (*green*) CFT073 bacteria plated on LB agar plates with MIC of amikacin or nitrofurantoin. Statistical significance was calculated using one-way ANOVA with a probability (*P*) value of <0.05 deemed statistically significant (\**P* <0.05, ns – not significant). Error bars represent standard deviation (n□= □3).

## Discussion

The σ subunits of the bacterial RNAP are central regulators of gene expression as they confer promoter-specificity upon the RNAP and thereby determine the expression of different regulons. The σ^54^ factor is distinct from other bacterial σ factors as it has an obligatory requirement for an activator ATPase to allow transcription by its RNAP and thus different activator ATPases determine the cognate regulon specificity of the σ^54^-RNAP in response to diverse environmental cues. This study has revealed a role for σ^54^ in the UPEC strain CFT073 as a determinant for homogeneous non-planktonic growth and, consequently, the absence of σ^54^ results in biphasic non-planktonic growth with two subpopulations appearing slower than the wild-type population on LB solid media. How the absence of σ^54^ affects the pathogenic capacity of UPEC bacteria remains to be investigated, but our results clearly indicate that the biphasic non-planktonic growth of the mutant bacteria imposes potentially adverse fitness consequences on the mutant population under growth restrictive conditions.

Although we are unable to specify the genetic basis by which σ^54^ determines homogeneous non-planktonic growth of the *E. coli* CFT073 strain, notably, the absence of σ^54^ results in the transcriptional upregulation of a subset of seemingly functionally unconnected genes. Intriguingly previous studies by Schaefer et al (21) and Shimada et al (22) implied that the σ^54^ or its RNAP could have a repressive effect on global transcription. It is thus possible that the transcriptionally attenuated RPc formed by the σ^54^-RNAP (see Introduction), when not required to be activated by the activator ATPase, could have an indirect regulatory function in controlling genetic information flow. Further, a study by Bonocora et al identified 85 intragenic binding events by σ^54^-RNAP mostly within σ^54^-independent genes in the genome of non-pathogenic *E. coli* strain MG1655 under a growth condition not associated with the activity of any activator ATPases, suggesting that most, if not all, of these promiscuous binding events could contribute to the transcriptional programme differently (23). Our results unequivocally indicate that σ^54^’s role as a determinant of homogeneous non-planktonic growth is not dependent on any of the individual activator ATPases present in the *E. coli* CFT073 strain. This is unsurprising as, to the best of our knowledge, exponential growth of *E. coli* in LB liquid media or LB solid media, does not require σ^54^. Hence, we propose that σ^54^’s role as a determinant of homogeneous non-planktonic growth represents a non-canonical function of σ^54^ in regulation of gene expression in bacteria. As σ^54^ is the only alternative σ factor that is abundant in *E. coli* during the exponential phase of growth in LB liquid media (i.e., the condition used here) (24), we envisage that σ^54^ functions like a nucleoid-associated protein (NAP) and thereby represses the transcription of genes required for homogeneous non-planktonic growth by the *E. coli* CFT073 strain. The observation that the absence of σ^54^ also results in the overall transcriptional upregulation of genes in the *E. coli* EDL933 strain, albeit not resulting in any detectable effect on non-planktonic growth, further supports a potential NAP-like function for σ^54^ in transcription regulation. Notably, different genes become transcriptionally upregulated in the Δ*rpoN* CFT073 and Δ*rpoN* EDL933 bacteria, suggesting that the non-canonical regulatory capacity of σ^54^ varies between strains of the same species. This might also explain why the absence of σ^54^ in either the EDL933 or NCM3722 *E. coli* strains does not result in biphasic non-planktonic growth on LB agar plates. Finally, in support of the potential NAP-like function for σ^54^, the Δ*rpoN* CFT073 strain harboring the plasmid pACYC-*rpoN* encoding for a σ^54^ variant containing the R456A substitution, which abrogates σ^54^’s ability to bind DNA but not RNAP (25), phenocopies the Δ*rpoN* CFT073 strain in the ScanLag assay (Fig. S1). It is difficult to decipher whether the potential NAP-like function of σ^54^ is mediated by σ^54^ *per se* or in the context of the RNAP. The former is a possibility as σ^54^, unlike any other six σ subunits in *E. coli*, can bind DNA independent of the RNAP (26).

Bacterial σ factors are considered as general transcription factors. In a recent review, Charles Dorman and colleagues proposed assigning a transcription regulatory protein as a ‘transcription factor’ or ‘NAP’ should be considered as *ad hoc* operational definitions (27). Hence, the potential NAP-like function of σ^54^ implied from our work further underscores this point and highlights the mechanistic complexity underpinning transcription regulation in bacteria.

## Materials and Methods

### Bacterial strains and plasmids

*Escherichia coli* strains and plasmid used in this study are listed in Table S1. The Δ*rpoN* mutants for all strains were made according to the λ Red recombinase method to replace the *rpoN* gene with an in-frame fusion encoding a kanamycin resistance cassette amplified from the pDOC-K plasmid (28). The Δ*rpoN*Δ*rpoS* CFT073 mutant strain was generated using the same recombinase method by replacing the *rpoS* gene with an in-frame fusion encoding a kanamycin resistant cassette using the Δ*rpoN* CFT073 strain as the parent strain. All mutant strains were cured of their kanamycin resistance cassette using the pCP20 plasmid (29). The *rpoN* complementation plasmid was constructed using Gibson assembly (30) to introduce the *rpoN* gene and ∼300 base pairs of the upstream native sequence (including the promoter) into a modified pACYC184 plasmid backbone (see supplementary Table S1 for details).

### Bacterial Growth

The planktonic growth assays were conducted in lysogeny broth (LB) liquid media or healthy female urine (taken from mid-stream flow) in flasks (25 ml) or 96-well plates at 37°C, shaking at ∼180 rpm. Growth curves were produced by measuring the optical density (OD_600nm_) of bacterial cultures as a function of time. All growth curves shown in figures represent the mean average from at least three biological replicates with the standard error of the mean (SEM) plotted as error bars. The doubling time in mid-log for growth curves was calculated according to Powell (31). Viability measurements were conducted by plating serial dilutions of cultures at relevant time points and counting colony forming units (c.f.u.) per ml of culture.

### Scanlag assays

Aliquots of bacterial cultures grown as described above were taken at time points indicated in the text, washed twice in sterile phosphate-buffered saline, diluted between 10^−5^ to 10^−6^, and 100 μl spread on either LB agar plates with or without specified antibiotics (2 μg/ml amikacin disulfate salt or 7 μg/ml nitrofurantoin (both Sigma-Aldrich) or 50% (v/v) healthy human urine containing 1.5% (w/v) agar. Plates were incubated at 33°C in a standard office scanner (Epson Perfection V370 photo scanner, J232D) placed in an incubator and images taken every 20 mins over a 48 h period. Analysis of appearance time and apparent growth rate of colonies was adapted from Levin-Reisman et al (17) using a modified code by Miles Priestman available at https://github.com/mountainpenguin/NQBMatlab. At least two independent experiments were conducted for each ScanLag data shown in the figures.

### RNA sequencing

Cultures were grown in LB liquid media and sampled during mid-exponential phase (OD_600nm_ = 1). Three biological replicates of each strain were taken and mixed with a phenol:ethanol (1:19) solution at a ratio of 9:1 (culture:solution) before harvesting the bacteria immediately by centrifugation. Bacterial pellets were sent to Vertis Biotechnologie AG for downstream processing. Briefly, RNA was extracted from the bacterial pellets using the RNAsnap (32) protocol followed by depletion of ribosomal RNA species, cDNA synthesis and next-generation sequencing with an Illumina NextSeq 500. Downstream data analysis was conducted using standard parameters on the CLC Genomics Workbench 7. RNA-seq reads for CFT073 and EDL933 genomes were mapped using Burrows-Wheeler Aligner to *E. coli* CFT073 (AE014075) and EDL933 (NZ_CP008957.1) strains, respectively. Unique reads mapped were used and total read counts used for data normalisation. Reads mapped to each gene were quantified to give a matrix of read counts, which was then analysed with the DESeq2 BioConductor package to identify differentially expressed genes. All statistical analysis for differential gene expression was conducted with R version 4.1.1. Sequencing data is available in the ArrayExpress database at EMBL-EBI (http://www.ebi.ack.uk/arrayexpress) under accession number E-MTAB-11288.

## Data Availability

Data will be shared upon request to the corresponding author, Sivaramesh Wigneshweraraj. *Author contributions:* A.S. and S.W. designed research; A.S. and L.B. performed research; A.S., P.M., and S.W. analysed data, and A.S. and S.W. wrote the paper.

## Funding and additional information

This work was supported by Wellcome Trust Investigator Award 100958 and Leverhulme Trust project grant RPG-2017-431.

## Conflict of interest

The authors declare that they have no conflicts of interest with the contents of this article.

### Abbreviations

The abbreviations used are:
RNAP: RNA polymerase
σ factor: sigma factor
σ^38^: RNAP promoted specificity factor S
σ^54^: RNAP promoted specificity factor N
σ^70^: RNAP promoted specificity factor D
bp: base pairs
RPc: closed promoter complex
RPo: open promoter complex
EHEC: enterrohemorrhagic *E. coli*
UPEC: uropathogenic *E. coli*
UTIs: urinary tract infections
IBCs: intracellular bacterial communities
h: hours
px^2^/h: pixels^2^ per hour
HFU: healthy female urine
v/v: volume per volume
MIC: minimum inhibitory concentration
NAP: nucleoid-associated protein
SEM: standard error of the mean
w/v: weight per volume
c.f.u.: colony forming units
ANOVA: analysis of variance
ns: not significant

## Supplementary material

**Figure S1.**
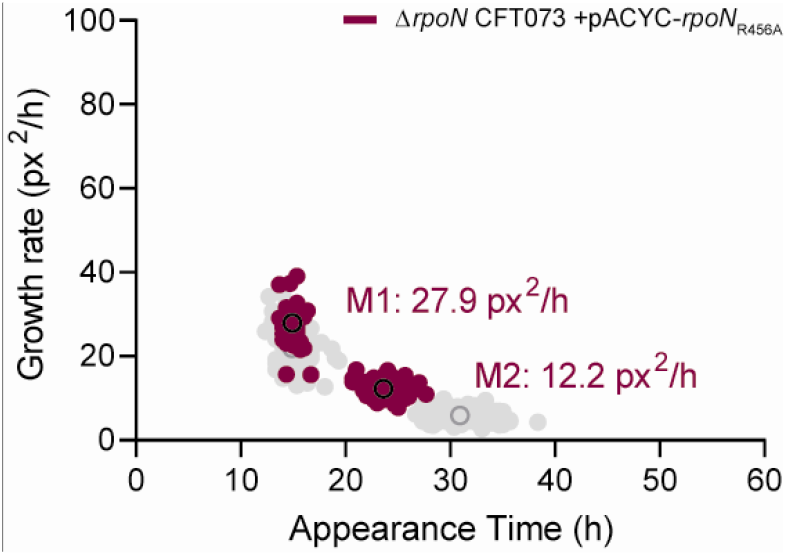
Scanlag analysis of colony appearance time following growth in LB liquid media for 5 h followed by plating on LB agar plates for Δ*rpoN* + pACYC-*rpoN*_R456A_ CFT073 bacteria. Black circles represent average population growth rate and appearance time with mean growth rate indicated as a value. Shown in grey for comparison are the M1 and M2 colonies from Δ*rpoN* CFT073 bacteria grown in LB liquid media for 5 h before plating.

**Table S1.** *E. coli* strains and plasmids used in this study

| Strains |  |  |
| --- | --- | --- |
| Name | Description | Source or Reference |
| Wild-type CFT073 | UPEC O6:K2:H1 WAM4505, original patient isolate | Gifted from Rodney Welch (1) |
| $\Delta rpoN$ CFT073 | UPEC O6:K2:H1 WAM4505, $\Delta rpoN$ | This study |
| $\Delta rpoN$ CFT073 + pACYC-*rpoN* | UPEC O6:K2:H1 WAM4505, $\Delta rpoN$ + pACYC-*rpoN* | This study |
| Wild-type EDL933 | EHEC O157:H7 strain, EDL933 | (2) |
| $\Delta rpoN$ EDL933 | EHEC O157:H7 strain, EDL933 $\Delta rpoN$ | This study |
| $\Delta rpoN$ EDL933 + pACYC-*rpoN* | EHEC O157:H7 strain, EDL933 $\Delta rpoN$ + pACYC-*rpoN* | This study |
| Wild-type NCM3722 | Prototrophic K-12 strain, NCM3722 | (3) |
| $\Delta rpoN$ NCM3722 | Prototrophic K-12 strain, NCM3722 $\Delta rpoN$ | This study |
| $\Delta rpoN$ NCM3722 + pACYC-*rpoN* | Prototrophic K-12 strain, NCM3722 $\Delta rpoN$ + pACYC-*rpoN* | This study |
| Plasmids |  |  |
| Name | Description | Source or reference |
| pACYC-*rpoN* | Modified pACYC plasmid backbone (-TcR, +MCS) expressing *rpoN* under the native *rpoN* promoter | This study |

**Table S2.**
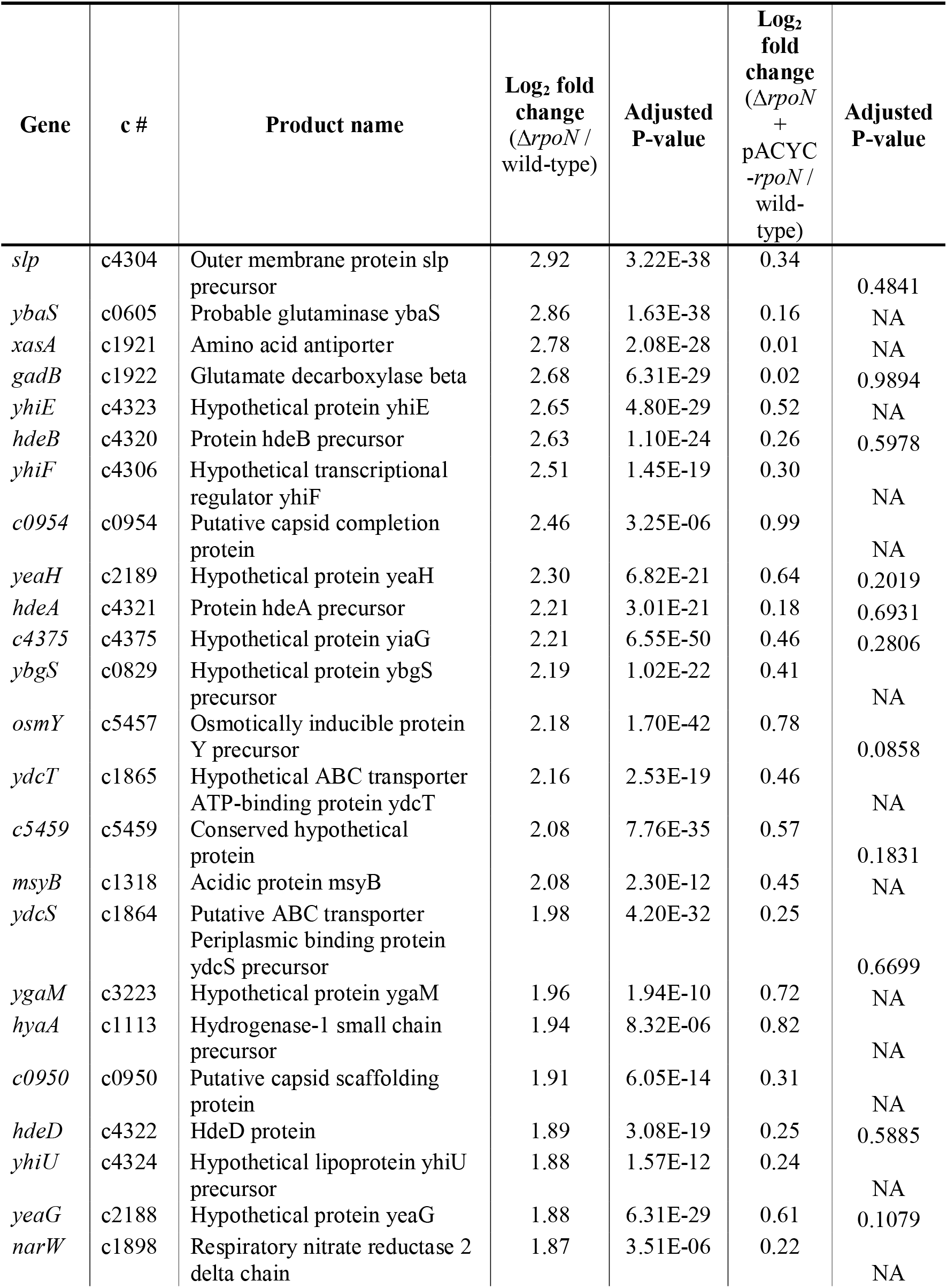
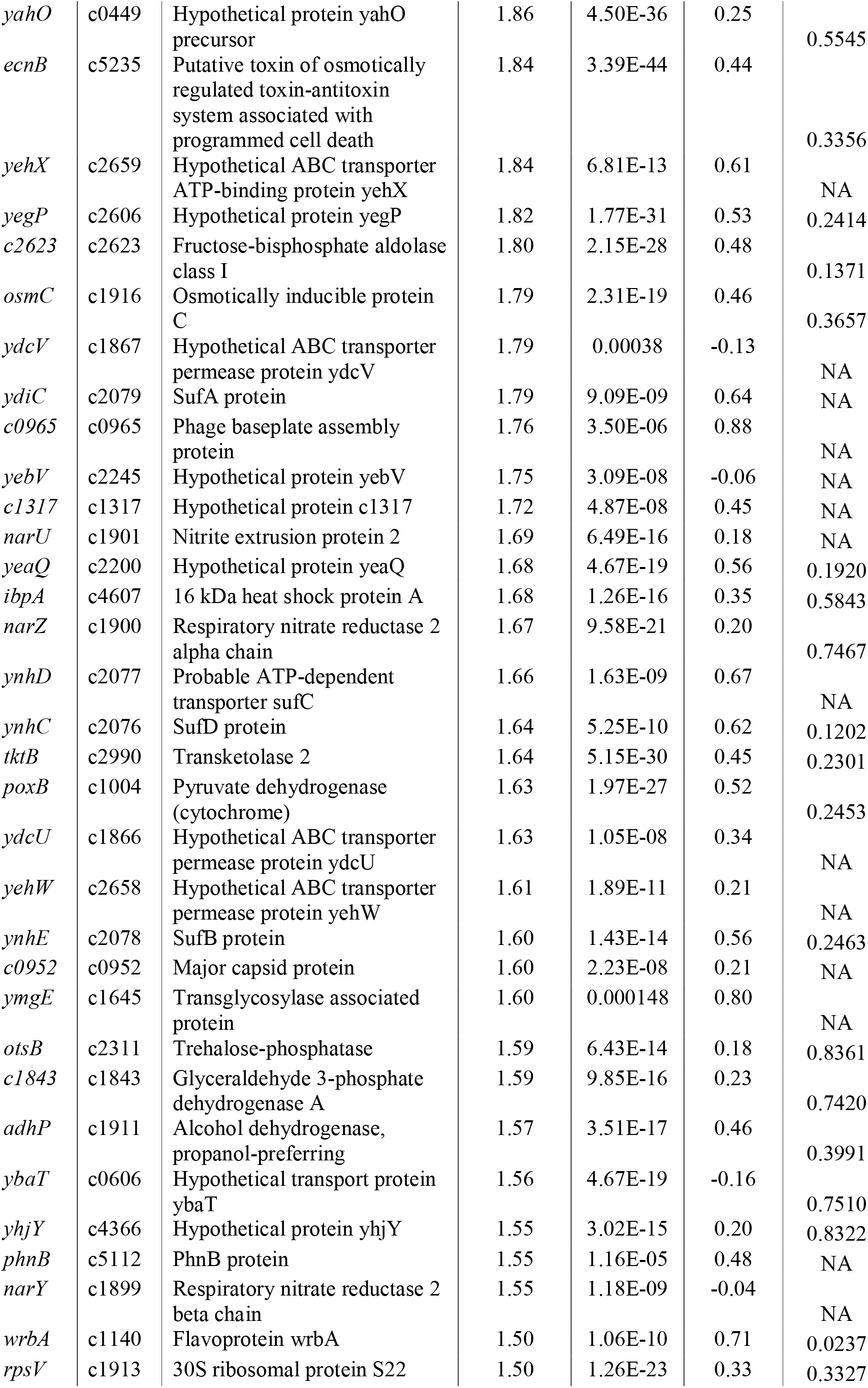
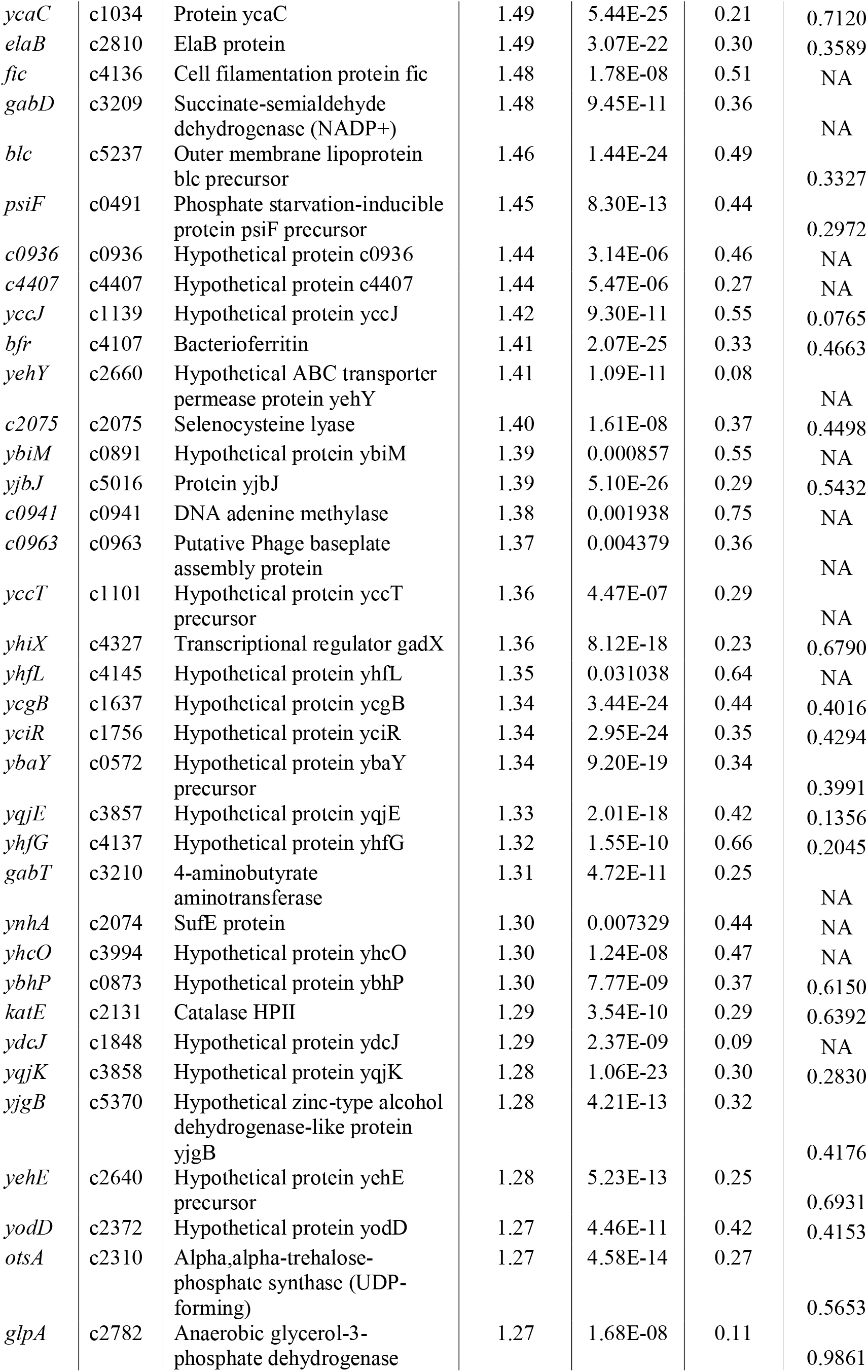
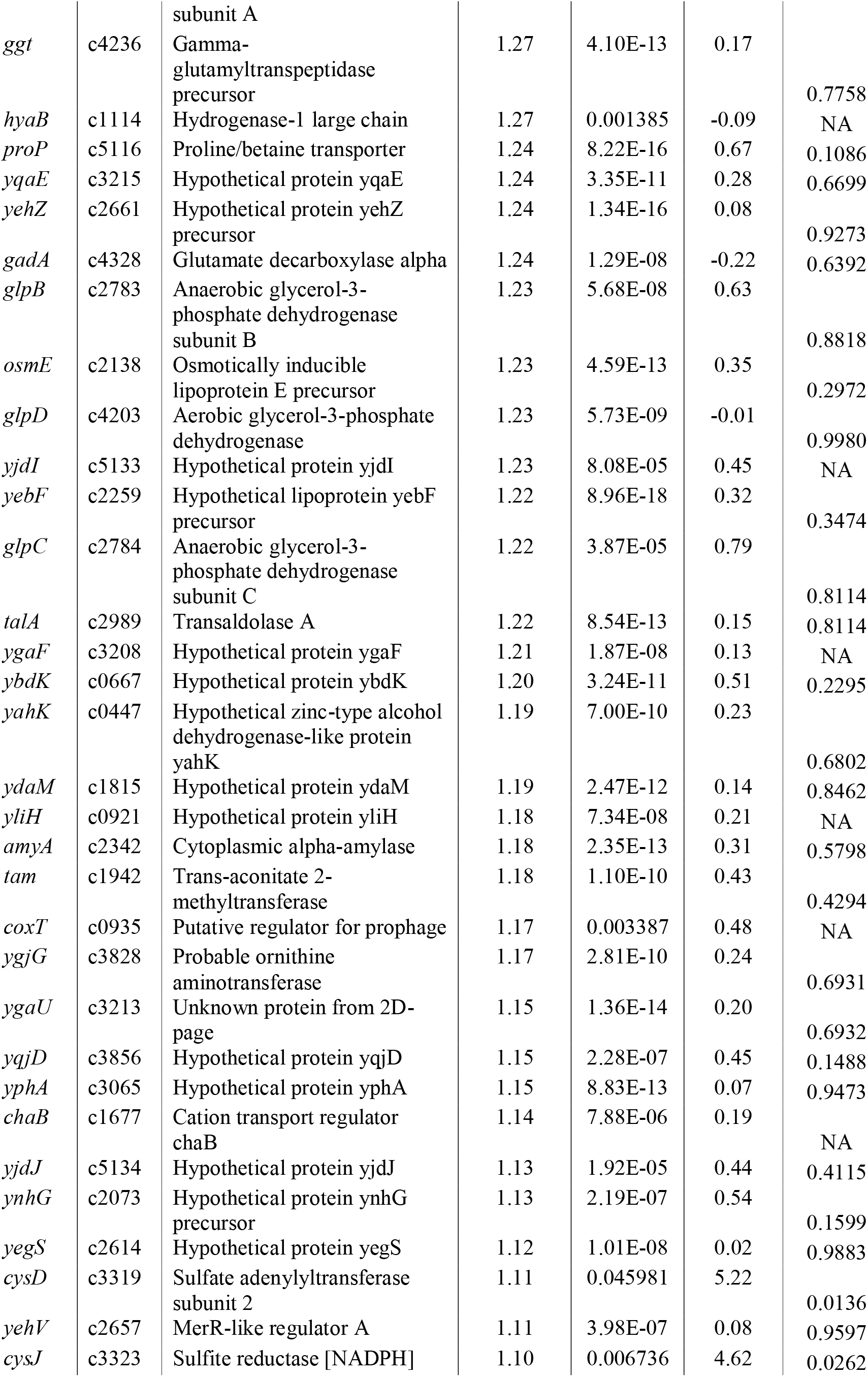
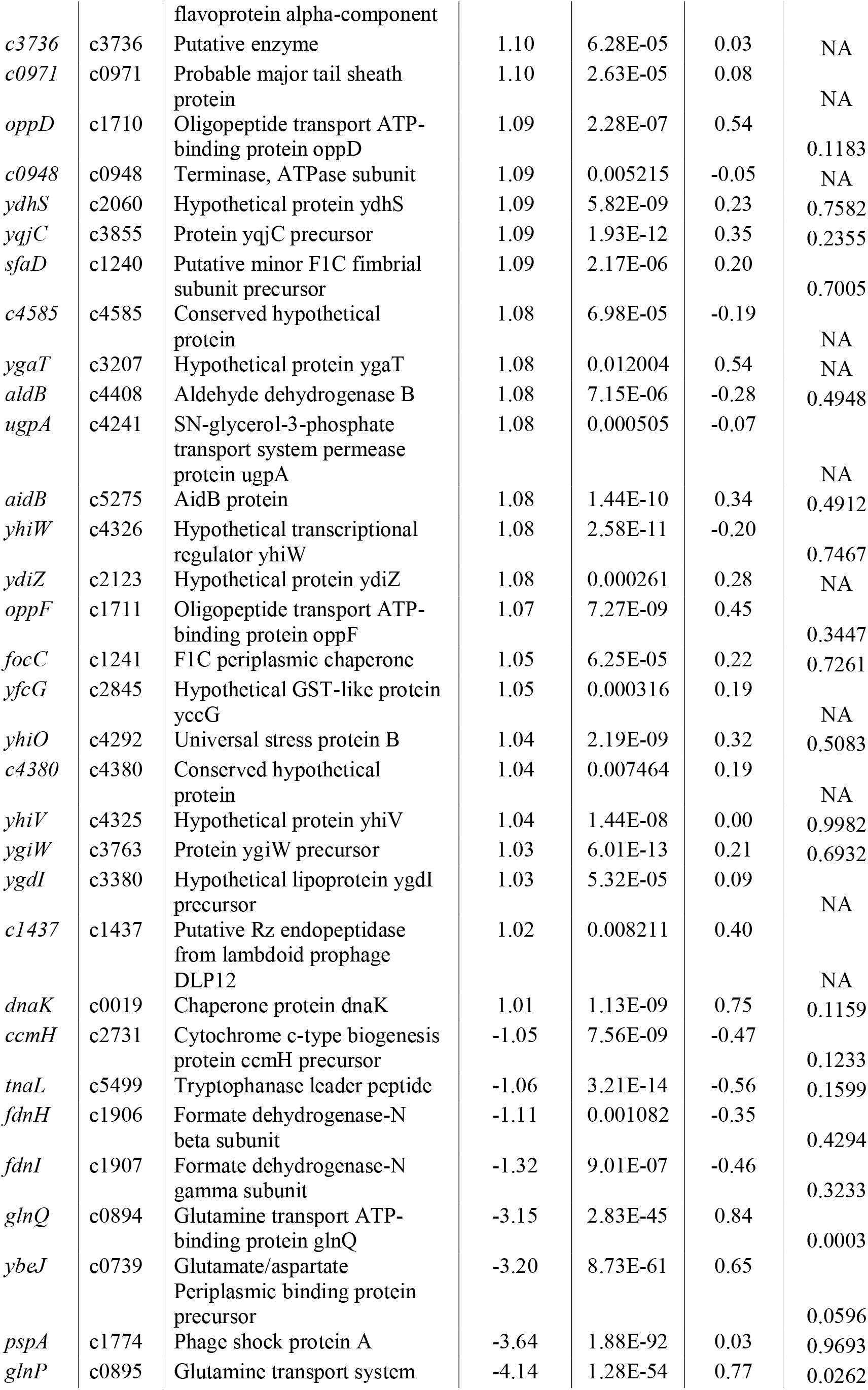
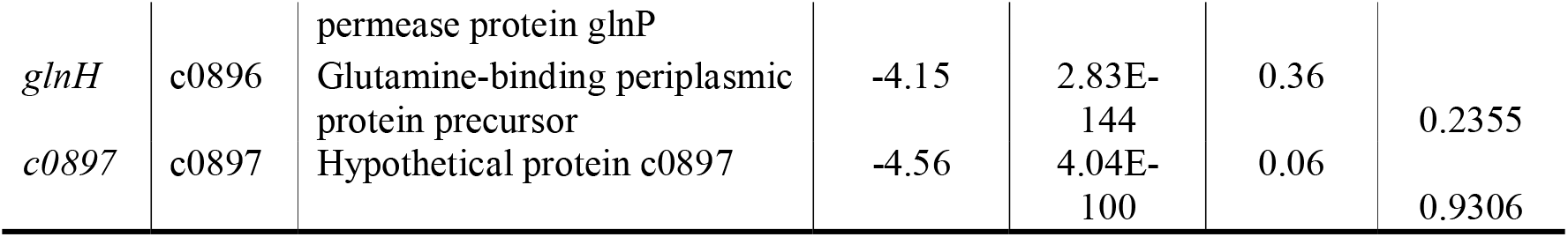
Differentially expressed genes in Δ*rpoN* CFT073 bacteria relative to wild-type CFT073 bacteria

**Table S3.**
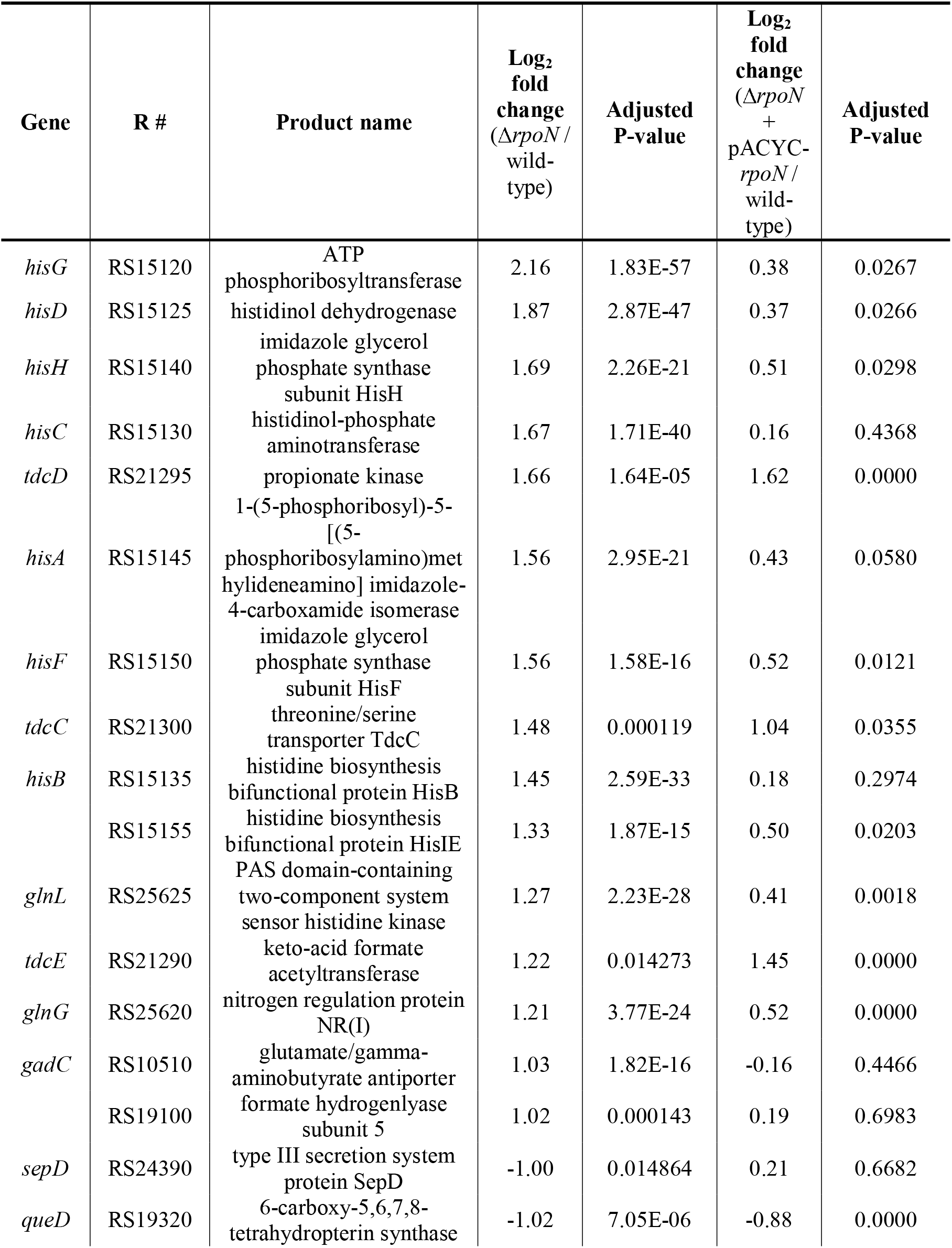
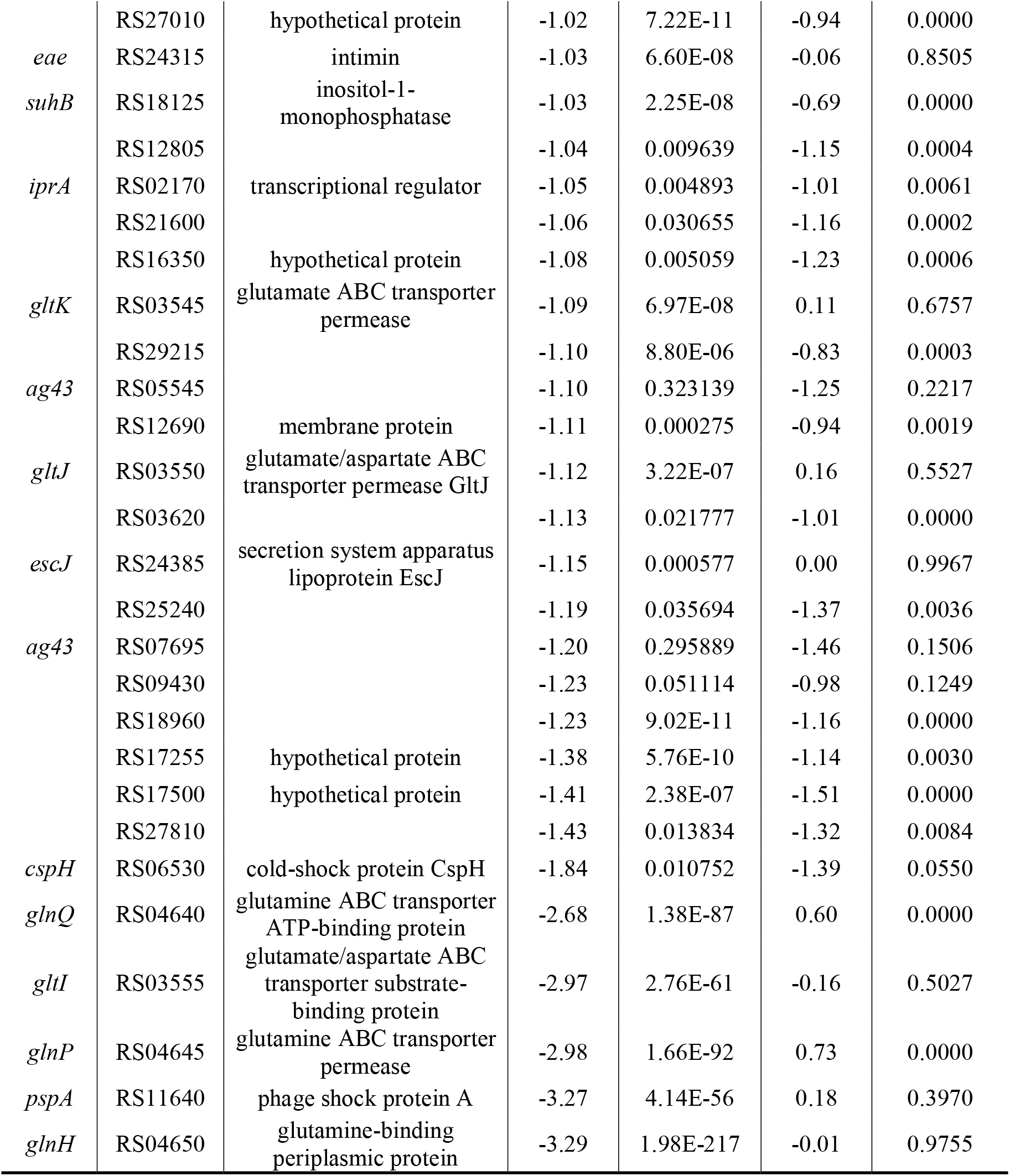
Differentially expressed genes in Δ*rpoN* EDL933 bacteria relative to wild-type and Δ*rpoN* + pACYC-*rpoN* EDL933 bacteria

| Gene | R # | Product name | Log<sub>2</sub> fold change ($\Delta rpoN$ / wild-type) | Adjusted P-value | Log<sub>2</sub> fold change ($\Delta rpoN$ + pACYC-*rpoN* / wild-type) | Adjusted P-value |
| --- | --- | --- | --- | --- | --- | --- |
| *hisG* | RS15120 | ATP phosphoribosyltransferase | 2.16 | 1.83E-57 | 0.38 | 0.0267 |
| *hisD* | RS15125 | histidinol dehydrogenase | 1.87 | 2.87E-47 | 0.37 | 0.0266 |
| *hisH* | RS15140 | imidazole glycerol phosphate synthase subunit HisH | 1.69 | 2.26E-21 | 0.51 | 0.0298 |
| *hisC* | RS15130 | histidinol-phosphate aminotransferase | 1.67 | 1.71E-40 | 0.16 | 0.4368 |
| *tdcD* | RS21295 | propionate kinase | 1.66 | 1.64E-05 | 1.62 | 0.0000 |
| *hisA* | RS15145 | 1-(5-phosphoribosyl)-5-[(5-phosphoribosylamino)met hylideneamino] imidazole-4-carboxamide isomerase | 1.56 | 2.95E-21 | 0.43 | 0.0580 |
| *hisF* | RS15150 | imidazole glycerol phosphate synthase subunit HisF | 1.56 | 1.58E-16 | 0.52 | 0.0121 |
| *tdcC* | RS21300 | threonine/serine transporter TdcC | 1.48 | 0.000119 | 1.04 | 0.0355 |
| *hisB* | RS15135 | histidine biosynthesis bifunctional protein HisB | 1.45 | 2.59E-33 | 0.18 | 0.2974 |
|  | RS15155 | histidine biosynthesis bifunctional protein HisIE | 1.33 | 1.87E-15 | 0.50 | 0.0203 |
| *glnL* | RS25625 | PAS domain-containing two-component system sensor histidine kinase | 1.27 | 2.23E-28 | 0.41 | 0.0018 |
| *tdcE* | RS21290 | keto-acid formate acetyltransferase | 1.22 | 0.014273 | 1.45 | 0.0000 |
| *glnG* | RS25620 | nitrogen regulation protein NR(I) | 1.21 | 3.77E-24 | 0.52 | 0.0000 |
| *gadC* | RS10510 | glutamate/gamma-aminobutyrate antiporter | 1.03 | 1.82E-16 | -0.16 | 0.4466 |
|  | RS19100 | formate hydrogenlyase subunit 5 | 1.02 | 0.000143 | 0.19 | 0.6983 |
| *sepD* | RS24390 | type III secretion system protein SepD | -1.00 | 0.014864 | 0.21 | 0.6682 |
| *queD* | RS19320 | 6-carboxy-5,6,7,8-tetrahydropterin synthase | -1.02 | 7.05E-06 | -0.88 | 0.0000 |
|  | RS27010 | hypothetical protein | -1.02 | 7.22E-11 | -0.94 | 0.0000 |
| *eae* | RS24315 | intimin | -1.03 | 6.60E-08 | -0.06 | 0.8505 |
| *suhB* | RS18125 | inositol-1-monophosphatase | -1.03 | 2.25E-08 | -0.69 | 0.0000 |
|  | RS12805 |  | -1.04 | 0.009639 | -1.15 | 0.0004 |
| *iprA* | RS02170 | transcriptional regulator | -1.05 | 0.004893 | -1.01 | 0.0061 |
|  | RS21600 |  | -1.06 | 0.030655 | -1.16 | 0.0002 |
|  | RS16350 | hypothetical protein | -1.08 | 0.005059 | -1.23 | 0.0006 |
| *gltK* | RS03545 | glutamate ABC transporter permease | -1.09 | 6.97E-08 | 0.11 | 0.6757 |
|  | RS29215 |  | -1.10 | 8.80E-06 | -0.83 | 0.0003 |
| *ag43* | RS05545 |  | -1.10 | 0.323139 | -1.25 | 0.2217 |
|  | RS12690 | membrane protein | -1.11 | 0.000275 | -0.94 | 0.0019 |
| *gltJ* | RS03550 | glutamate/aspartate ABC transporter permease GltJ | -1.12 | 3.22E-07 | 0.16 | 0.5527 |
|  | RS03620 |  | -1.13 | 0.021777 | -1.01 | 0.0000 |
| *escJ* | RS24385 | secretion system apparatus lipoprotein EscJ | -1.15 | 0.000577 | 0.00 | 0.9967 |
|  | RS25240 |  | -1.19 | 0.035694 | -1.37 | 0.0036 |
| *ag43* | RS07695 |  | -1.20 | 0.295889 | -1.46 | 0.1506 |
|  | RS09430 |  | -1.23 | 0.051114 | -0.98 | 0.1249 |
|  | RS18960 |  | -1.23 | 9.02E-11 | -1.16 | 0.0000 |
|  | RS17255 | hypothetical protein | -1.38 | 5.76E-10 | -1.14 | 0.0030 |
|  | RS17500 | hypothetical protein | -1.41 | 2.38E-07 | -1.51 | 0.0000 |
|  | RS27810 |  | -1.43 | 0.013834 | -1.32 | 0.0084 |
| *cspH* | RS06530 | cold-shock protein CspH | -1.84 | 0.010752 | -1.39 | 0.0550 |
| *glnQ* | RS04640 | glutamine ABC transporter ATP-binding protein | -2.68 | 1.38E-87 | 0.60 | 0.0000 |
| *gltI* | RS03555 | glutamate/aspartate ABC transporter substrate-binding protein | -2.97 | 2.76E-61 | -0.16 | 0.5027 |
| *glnP* | RS04645 | glutamine ABC transporter permease | -2.98 | 1.66E-92 | 0.73 | 0.0000 |
| *pspA* | RS11640 | phage shock protein A | -3.27 | 4.14E-56 | 0.18 | 0.3970 |
| *glnH* | RS04650 | glutamine-binding periplasmic protein | -3.29 | 1.98E-217 | -0.01 | 0.9755 |

